# Identifying Neuropsychiatric Disorder Subtypes and Subtype-dependent Variation in Diagnostic Deep Learning Classifier Performance

**DOI:** 10.1101/2022.10.27.514124

**Authors:** Charles A. Ellis, Robyn L. Miller, Vince D. Calhoun

**Affiliations:** Tri-institutional Center for Translational Research in Neuroimaging and Data Science at Georgia State University, Emory University, and the Georgia Institute of Technology

**Keywords:** dynamic functional network connectivity, schizophrenia, subtyping, deep learning, clinical decision support systems

## Abstract

Clinicians and developers of deep learning-based neuroimaging clinical decision support systems (CDSS) need to know whether those systems will perform well for specific individuals. However, relatively few methods provide this capability. Identifying neuropsychiatric disorder subtypes for which CDSS may have varying performance could offer a solution. Dynamic functional network connectivity (dFNC) is often used to study disorders and develop neuroimaging classifiers. Unfortunately, few studies have identified neurological disorder subtypes using dFNC. In this study, we present a novel approach with which we identify 4 states of dFNC activity and 4 schizophrenia subtypes based on their time spent in each state. We also show how the performance of an explainable diagnostic deep learning classifier is subtype-dependent. We lastly examine how the dFNC features used by the classifier vary across subtypes. Our study provides a novel approach for subtyping disorders that (1) has implications for future scientific studies and (2) could lead to more reliable CDSS.

## 1. INTRODUCTION

If neuroimaging clinical decision support systems (CDSS) are ever to be implemented in a clinical setting, they must be both robust and reliable [1]. One aspect of this reliability is that clinicians and need to not only know whether there are systematic differences in how the model will perform for different patients [2]. Neurological and neuropsychological disorder subtyping could contribute to more reliable CDSS [3][4]. While also providing scientific insights, the identification of subtypes could help identify patients for whom decision support systems will not perform well [3], thus helping reduce the likelihood of mistaken recommendations that might lead to misdiagnosis or incorrect treatment. In this study, we present a novel sequential clustering approach for subtyping neurological and neuropsychiatric disorders with neuroimaging data and evaluating diagnostic classifier performance on those subtypes.

While there are many different approaches for neuroimaging analysis, one common approach for the analysis of resting state functional magnetic resonance imaging (rs-fMRI) data involves using functional network connectivity (FNC). There are two types of FNC: static (sFNC, i.e., the average correlation between brain networks across a whole recording) and dynamic (dFNC, i.e., the correlation between brain networks in short increments over time). Both sFNC and dFNC have been used extensively to study a variety of neurological and neuropsychiatric disorders [5], [6] and cognitive capabilities [6], [7]. These analyses often involve clustering. The clustering of sFNC values has been used to identify subtypes or subgroups [7]. In contrast, dFNC values have mainly been used to identify neurological states but less so for subgroups or subtypes [5], [6]. That is unfortunate given that dFNC contains richer information than sFNC [8] and has routinely been used to identify links between neurological disorders and brain activity.

Separating dFNC into subtypes represents an important next step towards the eventual development of dFNC-based neuroimaging CDSS, as many proposed approaches for auditing CDSS involve the examination of differences in performance across subtypes [3][4]. Most classification studies using dFNC and sFNC have given insight into models performance for individual classes [9][10] but not subtypes. Discerning from the data of a patient whether a CDSS is likely to be accurate for them will be vital to ensuring that they receive proper care [2].

In this study, we present a novel sequential clustering approach for the identification of schizophrenia subtypes from dFNC data. We then develop an explainable 1-dimensional convolutional neural network (1D-CNN) to classify individuals with schizophrenia (SZs) and healthy controls (HCs). Lastly, we show how our model performance varies across subtypes and highlight the brain networks important for classifying each subtype.

## 2. METHODS

### 2.1. Description of Dataset

We used the Functional Imaging Biomedical Informatics Research Network (FBIRN) rs-fMRI dataset as a proof of concept. It has been used in many FNC studies [11][12]. The data can be made available upon a request to the authors. It was collected from the University of North Carolina at Chapel Hill, the University of California at Irvine, the University of New Mexico, the University of California at Los Angeles, Duke University, the University of California at San Francisco, the University of Iowa, and the University of Minnesota. The dataset consists of recordings from 151 individuals with schizophrenia (SZs) and 160 healthy controls (HCs). All study participants gave written informed consent, and data collection was approved by the Institutional Review Boards of the various institutions. Six 3T Siemens scanners and one 3T General Electric scanner were used during collection. T2*-weighted functional images were collected with an AC-PC aligned echo-planar imaging (EPI) sequence with TE=30ms, TR=2s, voxel size=3.4×3.4×4mm^3^, flip angle=77°, slice gap=1mm, 162 frames, and 5:24 minutes.

### 2.2. Description of Preprocessing and Feature Extraction

Preprocessing was conducted with statistical parametric mapping (https://www.fil.ion.ucl.ac.uk/spm/), and rigid body motion correction was used to account for head motion. Spatial normalization to an EPI template in the standard Montreal Neurological Institute space was performed. Data was resampled to 3×3×3 mm^3^ and smoothed with a Gaussian kernel (6 mm full width at half maximum). After preprocessing, we used the Neuromark automatic independent component (IC) analysis pipeline from the GIFT toolbox with the Neuromark_fMRI_1.0 template which has 53 ICs. The ICs were assigned to 7 networks. There were 5 subcortical (SCN), 2 auditory (ADN), 9 sensorimotor (SMN), 9 visual (VSN), 17 cognitive control (CCN), 7 default mode (DMN), and 4 cerebellar (CBN) ICs. To extract dFNC, we used a tapered sliding window where Pearson’s correlation was calculated between ICs at each step. The tapered window was a rectangle (40s window size) convolved with a Gaussian (standard deviation of 3). For each participant, this resulted in a 1378 feature x 124 time point matrix. The features are separable into 28 network pairs (e.g., SCN/ADN).

### 2.3. Description of Subtyping Approach

For SZ subtyping, we concatenated all data matrices across SZs to form a matrix of dimensions equal to all time steps across participants x dFNC features. We then performed k-means clustering, sweeping from 2 to 10 clusters, and selected the optimal number of clusters with the elbow method. The identified clusters represented states of dFNC activity that participants transitioned in and out of over time. We next calculated Occupancy Rates (OCR, i.e., percent of time spent in each state) for each participant and performed a second round of k-means clustering on the OCRs, sweeping from 2 to 10 clusters, and using the elbow method to select the optimal number of clusters. The clusters identified in this round of clustering represent data-driven subtypes of SZ. Previous studies have clustered the dFNC of combined SZ and HC participants to identify dFNC states [13][14], and some studies have calculated OCRs [12]. However, to our knowledge, no studies have used OCRs to further identify neurological disorder subtypes.

### 2.3. Description of Model Development

After subtyping SZs, we applied a 1D-CNN architecture from [9], [15]. We performed subject-specific, feature-wise z-scoring across all 124 time points. We used 10-fold stratified shuffle split cross-validation with an approximately 80-10-10 training-validation-testing split. We tripled the training set size via data augmentation consisting of adding Gaussian noise (μ = 0, σ = 0.7) to two copies of the training data. To account for class imbalances, we used a class-weighted categorical cross-entropy loss function. We used an Adam optimizer with an adaptive learning rate that started at 0.001 and decreased by 50% if 15 epochs passed without an increase in validation accuracy (ACC). We applied Kaiming He normal initialization and trained for 100 epochs with shuffling and a batch size of 50. We used model checkpoints to select the model from the epoch with the highest validation ACC for testing. To quantify performance, we calculated the mean and standard deviation of the ACC, sensitivity (SENS), and specificity (SPEC) across folds.

### 2.3. Description of Subtype Evaluation

To understand the interaction of the SZ subtypes with our model, we performed two analyses. (1) We looked for subtype-specific differences in model performance, and (2) we examined whether the model used different dFNC features for classifying each subtype.

To identify differences in subtype-specific performance, we calculated the model SENS for SZs belonging to each subtype in each fold. We then performed a Kruskal-Wallis nonparametric test to determine whether there were significant differences between model performance for subtypes across folds.

To determine whether the model relied upon different features for identifying each subtype, we used layer-wise relevance propagation (LRP). LRP has been used in a number of neuroscience time-series analyses [9], [15], [16]. We used the αβ-rule (α=1, β=0) to propagate positive relevance through the network (i.e., relevance that a sample belongs to a specific target class, rather than other classes). We applied LRP for all SZ test samples across folds, using SZ as the target class. We then normalized the relevance for each sample to sum to 100%, spatially summed the relevance across all time steps for each sample, averaged the LRP values for each subtype in each fold, and averaged for each subtype across folds.

## 3. RESULTS AND DISCUSSION

In this section, we describe and discuss our subtyping and model performance results. We also discuss study limitations and next steps.

### 3.1. Identification of 4 SZ Subtypes

We identified 4 dFNC states and 4 SZ subtypes. Our first round of clustering identified 4 dFNC states, shown in Figure 2. This finding is comparable to those of [13], [14], which identified 5 dFNC states for DMN-specific clustering [13] and clustering of whole brain dFNC across both SZs and HCs [13], [14]. State 3 has the highest magnitude correlations, followed by State 2. State 0 has very high intra-SCN, intra-SMN, intra-VSN, and intra-CBN activity. It also has high SMN/ADN, SMN/VSN, and VSN/ADN activity. State 3 also has very negative SCN/SMN, SCN/VSN, SMN/CBN, and VSN/CBN correlations. The remaining states have mainly positive correlations. State 2 has relatively high intra-SCN, intra-SMN, intra-VSN, and intra-CBN activity. States 0 and 1 have lower magnitude correlations, with State 0 have moderate intra-VSN and intra-SMN correlation and State 1 only having moderate intra-VSN activity.

**Figure 1.**
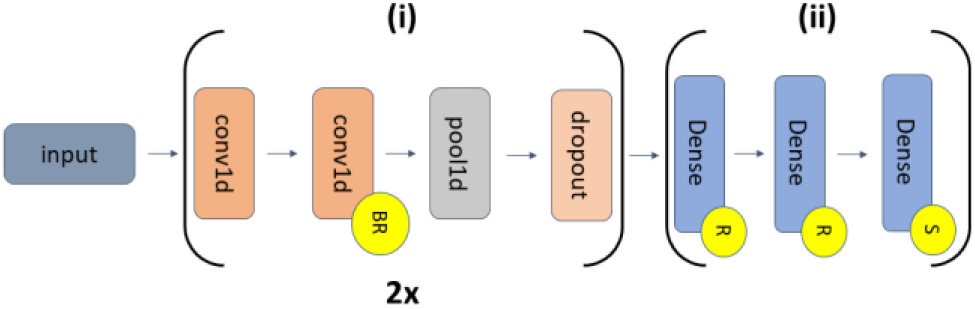
CNN Architecture. The model has feature extraction (i) and classification (ii) portions. (i) repeats twice. The first and second convolutional (conv1d) layer (kernel size = 10) pairs have 16 and 24 filters, respectively. Each pair is followed by a max pooling layer (pool size = 2) and spatial dropout (rates = 0.3 and 0.4). Portion (ii) has 3 dense layers with 10, 6, and 2 nodes, respectively. Yellow circles with “R”, “BR”, and “S” show layers with ReLU, batch normalization and ReLU, and softmax activations, respectively.

**Figure 2.**
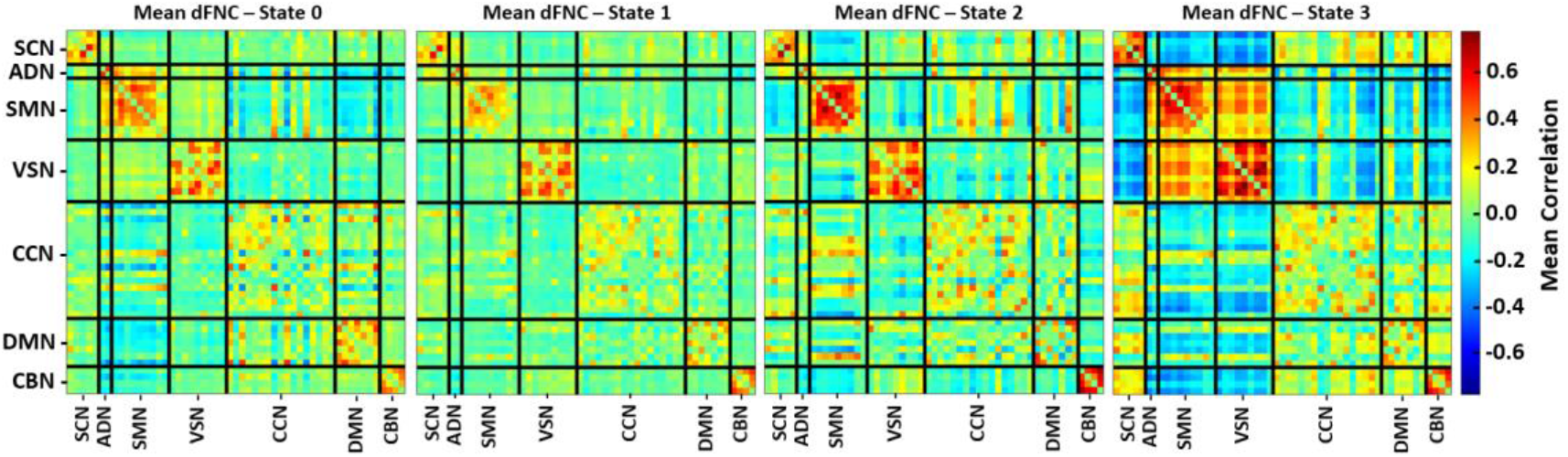
Mean dFNC for Each State. Each panel shows the mean dFNC (i.e., correlation between each of the 53 ICs) for a different state. The x- and y-axes indicate different networks into which the ICs are divided, and the black lines separate the connectivity of each network pair. All panels share the colorbar to the right. The magnitude of the connectivity between states varies greatly.

Our OCR clustering identified 4 SZ subtypes. Figure 3 shows the OCRs for each subtype. Subtypes 1, 2, 3, and 4 have 28, 56, 27, and 40 participants, respectively. Interestingly, subtype 1 is mainly state 3, subtype 2 is mainly state 1, subtype 3 is mainly state 2, and subtype 4 is mainly state 0. Nevertheless, many individuals in subtype 1 have non-zero state 0 OCRs, and the remaining subtypes spend some time in state 3. Additionally, many subtype 4 individuals have moderate state 1 OCRs. Subtype 1 spends time mainly in a highly connected brain state, while other subtypes spend only small amounts of time in a highly connected brain state. Subtype 2 spends more time in a state of moderate intra-VSN connectivity, and subtype 4 spends more time in a state of moderate intra-VSN and intra-SMN connectivity with less interaction between networks.

**Figure 3.**
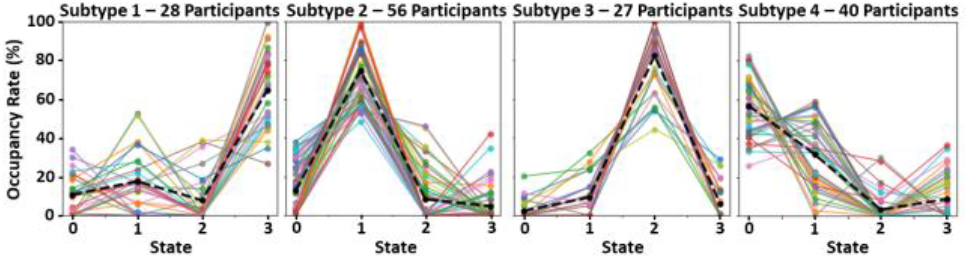
Subtype OCRs. Each panel shows the OCRs for a different subtype. The title of each panel indicates its subtype and the number of study participants in its subtype. The y-axis indicates the OCR, and the x-axis indicates each state. While each subtype spends some time in each state, each subtype spends more time in a different state.

### 3.2. Model Performance

Across folds, our model obtained a test SPEC, SENS, and ACC of 75.63 ± 14.64, 74.38 ± 11.24, and 75.00 ± 7.26, respectively. All performance metrics were well above chance-level. Model SPEC was higher than SENS. However, SPEC varied more across folds.

### 3.3. Subtype-Specific Differences in Model Performance

Figure 4 shows the differences in model performance for each subgroup across folds. Although our Kruskal-Wallis did identify significant differences in performance across folds, there were visible differences. The model generally performed highly for subtypes 2 and 4, which were larger and spent more time in poorly connected states. In contrast, its performance for subtypes 1 and 3 varied across folds, and subtype 1 had a much lower median value.

**Figure 4.**
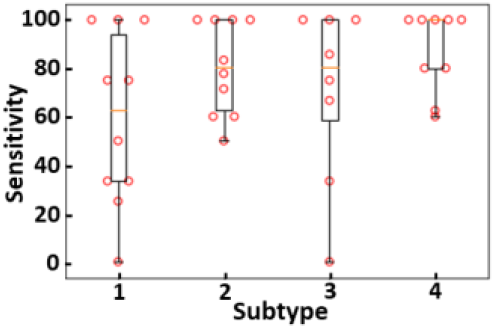
Subtype-specific Sensitivity. The y-axis shows the sensitivity of the model across folds, and the x-axis shows each subtype. Each circle indicates model sensitivity for a fold.The model performs well for subtypes 4 and varies in performance for the remaining subtypes.

### 3.4. Subtype-Specific Differences in Explanations

The mean LRP relevance values for each subgroup are shown in Figure 5. Interestingly, the relevance distributions tend to be fairly distinct across subtypes. For subtype 1, the model relied mainly upon the interaction of the SCN with other networks, which was the only subtype to spend time mostly in state 3 with highly negative SCN/SMN and SCN/VSN interactions. For subtypes 2 and 4, the model also relied upon CBN low connectivity with other networks (SCN, SMN) but to a lesser extent with subtype 4 than subtype 2. This could be why the model performed higher for those subtypes. Lastly, the model relied more upon VSN/SCN and VSN/SMN for subtype 3. These findings are interesting. In general, all subtypes tended to have some differences from HCs in their CBN/SCN and CBN/VSN or CBN/SMN. However, this was less so the case for subtypes 1 and 3, which might be considered SCN-and VSN-variant subtypes, respectively.

**Figure 5.**
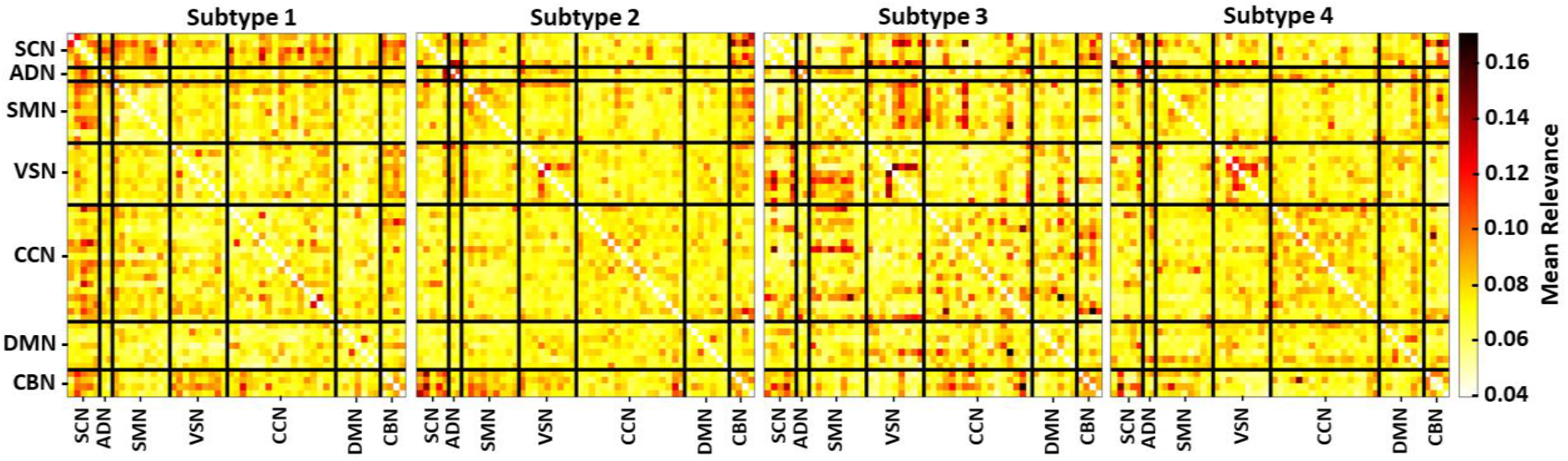
Mean Relevance for Each Subtype. Each panel shows the mean spatial relevance for a different subtype. The x- and y-axes indicate different networks into which the ICs are divided, and the black lines separate the connectivity of each network pair. All panels share the colorbar to the right. There are visible differences in distributions of relevance for each subtype.

### 3.5. Limitations and Next Steps

We only examined spatial differences in LRP relevance between groups. Future iterations might examine temporal differences in relevance between subtypes [9], [10]. Additionally, it might be helpful to apply a clustering explainability approach to quantify the relative importance of each network pair to each SZ subtype using a modified version of the methods used in existing studies [7], [11]. It would also be beneficial to examine the relationships between the identified dFNC subgroups and neuropsychological and clinical measures. Lastly, in this study, we obtained data-specific subtypes of SZ. It could also be useful to identify model-specific SZ subtypes.

## 4. CONCLUSION

Neurological and neuropsychiatric disorder subtyping could contribute to the development of more reliable deep learning-based neuroimaging CDSS by identifying specific patient groups for whom models may perform poorly. In this study, we present a novel subtyping approach for within the context of SZ using dFNC. After identifying 4 SZ subtypes, we show how the performance of a deep learning classifier trained for SZ diagnosis varies across subtypes and subsequently show how it relies upon different brain networks for identifying each subtype. Our method is broadly applicable for the subtyping of a variety of neurological and neuropsychiatric disorders and could eventually contribute to the development of more reliable deep learning-based neuroimaging CDSS.

## 5. ACKNOWLEDGMENTS

- We thank those who collected the FBIRN dataset.
- This work was supported by NSF 2112455 and NIH R01MH123610

